# RobMixReg: an R package for robust, flexible and high dimensional mixture regression

**DOI:** 10.1101/2020.08.02.233460

**Authors:** Wennan Chang, Changlin Wan, Chun Yu, Weixin Yao, Chi Zhang, Sha Cao

**Affiliations:** Department of Electrical and Computer Engineering, Purdue University, Indianapolis, IN, 46202, USA; Department of Medical and Molecular Genetics, Department of Biostatistics, Center for Computational Biology and Bioinformatics, Indiana University School of Medicine, Indianapolis, IN,46202, USA; School of Statistics, Jiangxi University of Finance and Economics, Nanchang, China; Department of Statistics, University of California, Riverside, CA, USA

## Abstract

**Motivation:** Mixture regression has been widely used as a statistical model to untangle the latent subgroups of the sample population. Traditional mixture regression faces challenges when dealing with: 1) outliers and versatile regression forms; and 2) the high dimensionality of the predictors. Here, we develop an R package called RobMixReg, which provides comprehensive solutions for robust, flexible as well as high dimensional mixture modeling.

**Availability and Implementation:** RobMixReg R package and associated documentation is available at CRAN: https://CRAN.R-project.org/package=RobMixReg.

## 1 Introduction

The big “Omics” data era has profoundly changed biology by scaling data acquisition, and transformed the biomedical research ecosystem from traditional case-based studies to modern large scale data driven research. For example in cancer research, the Cancer Genome Atlas (TCGA) has collected biospecimens and matched clinical phenotypes for over 10,000 cancer patients (Weinstein, et al., 2013). Driven by the increasing availability of phenotypic and multiple omics data, researchers have gained unprecedented opportunity to interrogate biology in novel and creative ways. Regression-based association analysis remains to be the most popular data mining tool for hypothesis generation, owing to its ease of interpretability. However, the traditional linear regression model cannot adequately fit the complexity of the biomedical data caused by: 1) high leverage noise and outliers resulted from the large scale technical experiments; 2) the presence of subpopulations among the wide spectrum of collected samples; 3) the high-dimensionality of the molecular features. This raises an urgent demand of a toolkit to specifically handle these challenges, and more importantly, evaluate the level of outlier contamination, the level of heterogeneity, as well as the level of sparseness of the features, or in other words, the most optimal model to fit the data.

We have developed an R package called “RobMixReg” to provide a comprehensive solution to mining the latent relationships in biomedical data. The major contribution of RobMixReg lies in the following key aspects: 1) for low dimensional predictors, RobMixReg allows for robust parameter estimations, and detection of the outliers with adaptive trimming; 2) RobMixReg allows different mixture components to have flexible forms of predictors; 3) RobMixReg handles the high-dimensional predictors by regularizing the regression coefficients of each component, with a data-driven level of sparsity; 4) RobMixReg provides the capability for order selection. To the best of our knowledge, there is no software package having these comprehensive functionalities and flexibility with similar purposes as RobMixReg in mining the biomedical data. We believe RobMixReg will be a valuable addition to the biomedical research community.

**Table 1.**
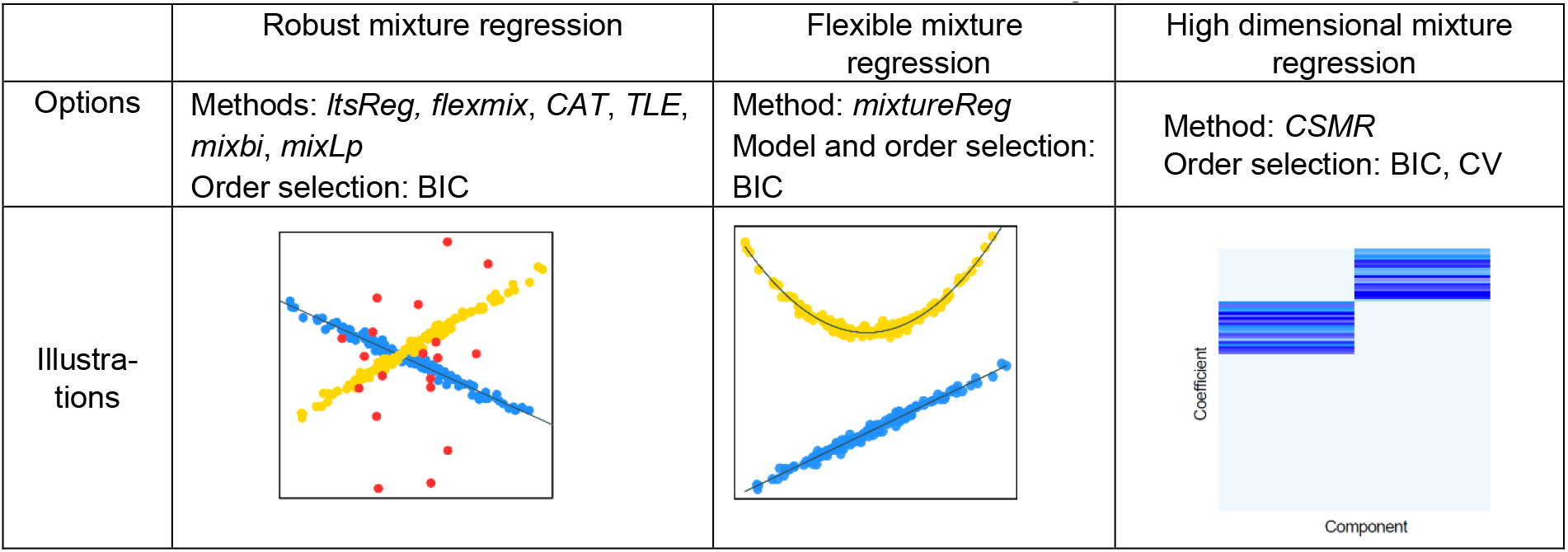
The functionalities of RobMixReg

|  | Robust mixture regression | Flexible mixture regression | High dimensional mixture regression |
| --- | --- | --- | --- |
| Options | Methods: *ItsReg*, *flexmix*, *CAT*, *TLE*, *mixbi*, *mixLp*<br>Order selection: BIC | Method: *mixtureReg*<br>Model and order selection: BIC | Method: *CSMR*<br>Order selection: BIC, CV |
| Illustrations |  |  |  |

## 2 Methods

Finite Mixture Gaussian Regression (FMGR) is a widely used model to explore the latent relationship between the response and predictors (Böhning, 1999). When fitting a *K*-component FMGR model with *N* observations of response variable *y* and *P*-variate predictor ***x***, we usually consider maximizing the following likelihood function:

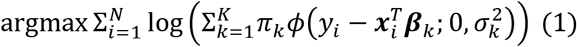

where *π_k_, β_k_, σ_k_* are the prior probability, the regression coefficients and the standard deviation of the *k*-th component; *ϕ*(·) is the normal density function. The unknown parameters in the above FMGR model is usually estimated by the EM algorithm (Dempster, et al., 1977),

RobMixReg deals with regression scenarios for both low and high dimensional predictors. As shown in Figure 1, for low dimensional predictors, RobMixReg considers robust and flexible mixture modeling; for high dimensional predictors, RobMixReg considers adaptive feature selection. In addition, for both cases, it provides a way for order selection. More details on implementation of these methods could be found in the supplementary document.

**Figure 1:**
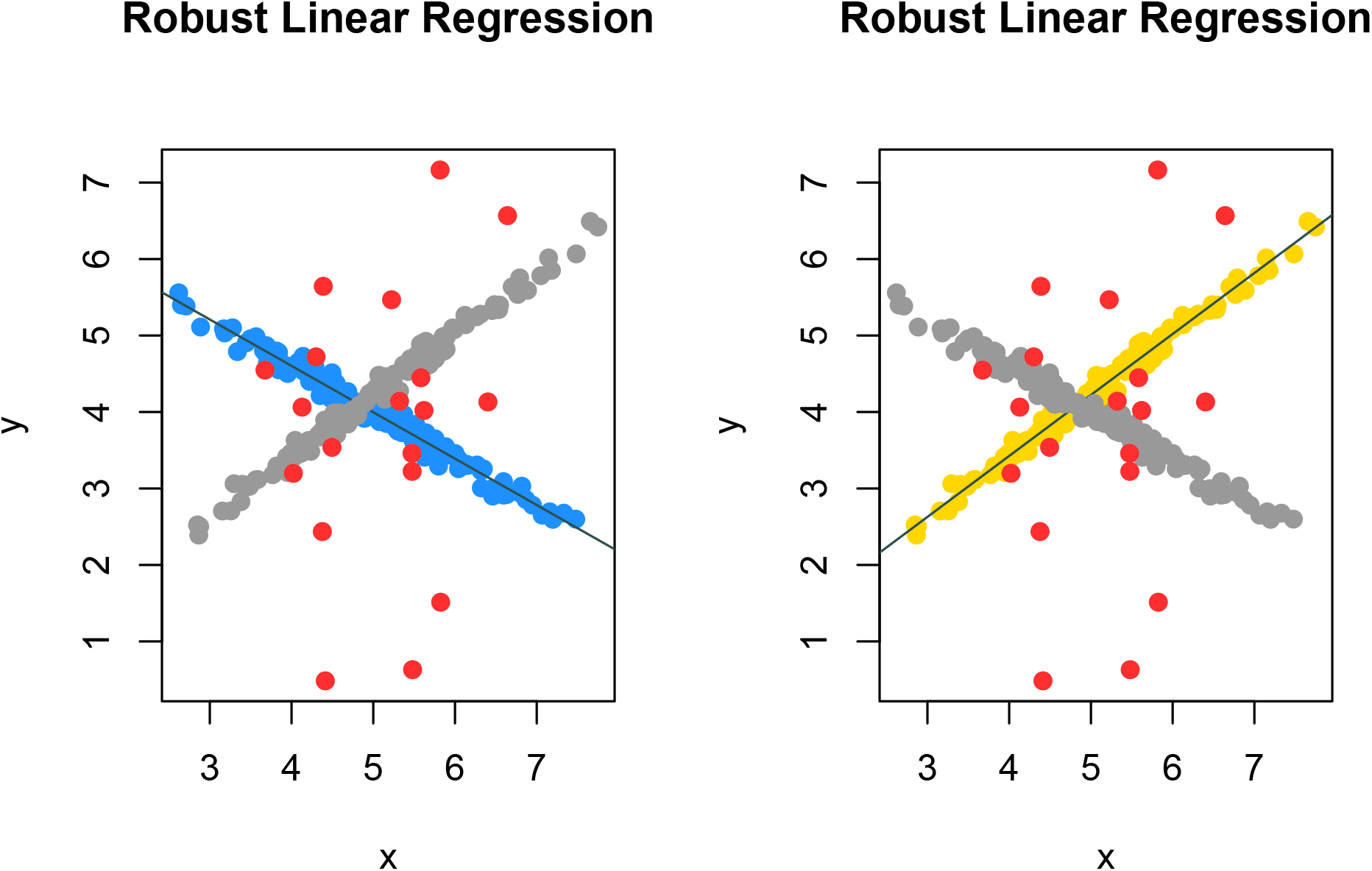
Robust mixture regression result

### 2.1 Robust mixture regression

In RobMixReg, the traditional EM algorithm was implemented in by the *flexmix* function in the *flexmix* package (Leisch, 2004). The traditional EM algorithm is very sensitive to outliers due to the least square criterion used in the M step. Several robust approaches were proposed to estimate the FMGR parameters, including the trimmed likelihood estimator (*TLE*) (Neykov, et al., 2007), the Component-wise Adaptive Trimming (*CAT*) algorithm (Chang, et al., 2020), the a robust bi-square criterion method (*mixbi*) (Bai, et al., 2012), the robust mixture of t-distributions (*mixt*) (Yao, et al., 2014), robust mixture of Laplace distribution (*mixLp*) (Song, et al., 2014), and a mean shift mixture regression model (*RM2*) (Yu, et al., 2017). In the RobMixReg package, for *K* = 1, the robust mixture regression degenerates to robust linear regression, and the function *ltsReg* in the package *robust-base* is implemented. For *K* > 1, methods including the *flexmix*, *TLE*, *CAT*, *mixbi*, and *mixLp* are all implemented.

### 2.2 Flexible mixture regression

When fitting two or more lines through the data, there could be different predictors and their functional forms in different lines. For example, two lines may involve different subsets of the predictors; or a line could include nonlinear transformations of the predictors to relax the “linearity” constraints. We provide the capability for flexible mixture regression models such that the user could specify the regression functions for different components. Different from traditional FMGR model, where the predictors consist of only themselves, in flexible mixture regression, users could specify the predictors and the transformations of the predictors. Such a flexibility is important as the complexity of the model could be well controlled. We include a flexible method in our package to enable users to decide what predictors to be used for each mixing component developed in (Zhou, 2017).

### 2.3 High dimensional mixture regression

While mixture regression is capable of handling the heterogeneous relationships, it does not work well in the case of high dimensional molecular features, where the total number of parameters to be estimated is far more than the total number of observations. Hence, restrictions on the sparsity level of ***β***_*k*_ needs to be introduced to shrink the insignificant regression coefficients. RobMixReg implemented the *CSMR* algorithm (Chang, et al., 2020), which performs simultaneous sample clustering and feature selection. It has a built-in functionality for the automatic selection of the sparsity level, which substantially improves both the computational efficiency and biological interpretability.

### 2.4 Order selection

Determining the order (number of components, *K*) for mixture models has long been a challenging and important problem. We use the Bayesian information criterion (BIC) for order selection, which is calculated as:

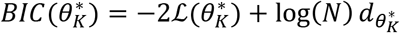

Here, 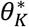 denotes the parameter estimates for component number *K*, and 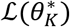 is the associated log likelihood; *N* is the total number of observations; 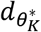 is the total number of parameters, and 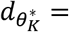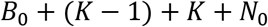. Here, *B*_0_ is the total number of non-zero regression coefficients; *K* − 1, the number of component proportions; *K*, the number of standard deviations for the *K* components; *N*_0_, the total number of outliers detected. Note that, we currently do not consider outliers for the flexible and high dimensional settings, hence, *N*_0_ = 0 in these cases. BIC criterion could be also used for model selection in flexible mixture regression. In addition to the BIC criteria, RobMixReg also offers a cross validation algorithm for the selection of *K* for the high dimensional case.

## 3 Results

We used four simulation and four real data examples to demonstrate the capability of RobMixReg (see supplementary file), in dealing with robust mixture regression, flexible mixture regression and high dimensional mixture regression. We provided the solution for order selection in each scenario.

## Supplementary Materials

### S1. Introduction

Regression-based association analysis remains to be the most popular data mining tool for hypothesis generation, owing to its ease of inter-pretability. However, the traditional linear regression model cannot adequately fit the complexity of the biomedical data caused by: 1) high leverage noise and outliers resulted from the large scale technical experiments; 2) the presence of subpopulations among the wide spectrum of collected samples; 3) the high-dimensionality of the molecular features. Finite Mixture Gaussian Regression (FMGR) is a widely used model to explore the latent relationship between the response and predictors, and the goal of mixture regresison is to maximize the following likelihood:

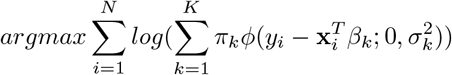

Here, we are modeling the relationship between **x** and *y* as a mixture of *K* regression lines, *K* ≥ 1. (**x**_**i**_, *y*_*i*_) is the *i*-th observations *i* = 1*, …, N*, *β*_*k*_ are the regression coefficients for the *k*-th regression line; *ϵ*_*ik*_ is the error term; *σ*_*k*_ is the standard deviation of *ϵ*_*ik*_.

‘RobMixReg’ is a package that provides a comprehensive solution to mining the latent relationships in biomedical data, ranging from the regulatory relationships between different molecular elements, to the the relationships between pheno-types and omic features, in the presence of outliers, latent subgroups and high dimensional molecular features.

The major contribution of RobMixReg lies in the following key aspects: 1) for low dimensional predictors, it handles outliers in modeling the latent relationships, and provides flexible modeling to allow for different predictors for different mixture components; 2) for high dimensional predictors, it detects subgroups and select informative features simultaneously. The RobMixReg package is a powerful exploratory tool to mining the latent relationships in data, which is often obfuscated by high leverage outliers, distinct subgroups and high dimensional features.

The RobMixReg package could be installed from CRAN:

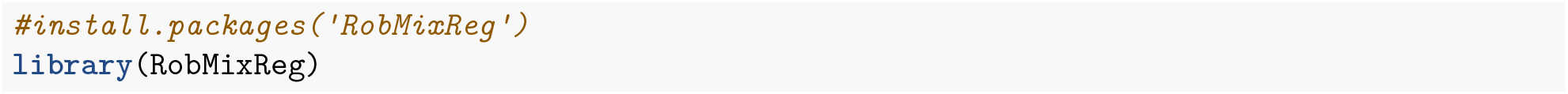

It could be also installed from GitHub:

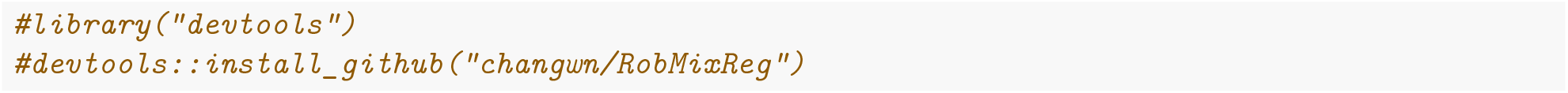

### S2. Case Studies

#### S2.0 Data Simulation

The function *simu_func* is used to generate the synthetic data for various of cases. The input of *simu_func* is:

- *β* A matrix whose *k*-th column is the regression coefficients matrix for the *k*-th component
- *σ* A vector whose *k*-th element is the standard deviation of the error term for the *k*-th regression component
- *alpha* The proportion of the observations being contaminated by the outlier

The output of *simu_func* is:

- *x* Matrix of the predictors with dimension *N* × (*P* + 1), where *P* is the total number of predictors, and the first column is an all-one vector.
- *y* Vector of response variable where *alpha* proportion of the observations are contaminated by outliers.

#### S2.1 Robust mixture regression

We adopt the mean-shift model in the presence of outliers, such that each outlier requires one additional parameter to cancel out the unusually large residuals. The outlier parameters are then regularized to achieve a balance of model complexity and model fitness.

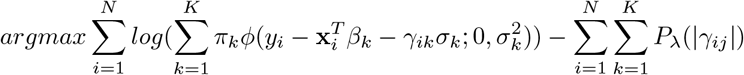

Here, *γ*_*ik*_ models the level of outlierness for the *i*-th observation; *P*_*λ*_(|*γ*_*ij*_|) is a penalty function with a tuning parameter *λ* controlling the degrees of penalization on the *γ*_*ik*_.

We have intergrated multiple state-of-the-art methods in our package to robustly fit two or more regression lines. The default method is CAT. Other options inlcude TLE, flexmix, mixLp, mixbi, and they could be passsed to the MLM function by the parameter ‘rmr.method’. Among them, CAT and TLE could detect the identities of the outliers, and outliers are removed before model estimation; while the rest of the methods perform model estimation in the presence of outliers. For TLE to work, a trimming proportion needs to be provided, and the default is 0.05.

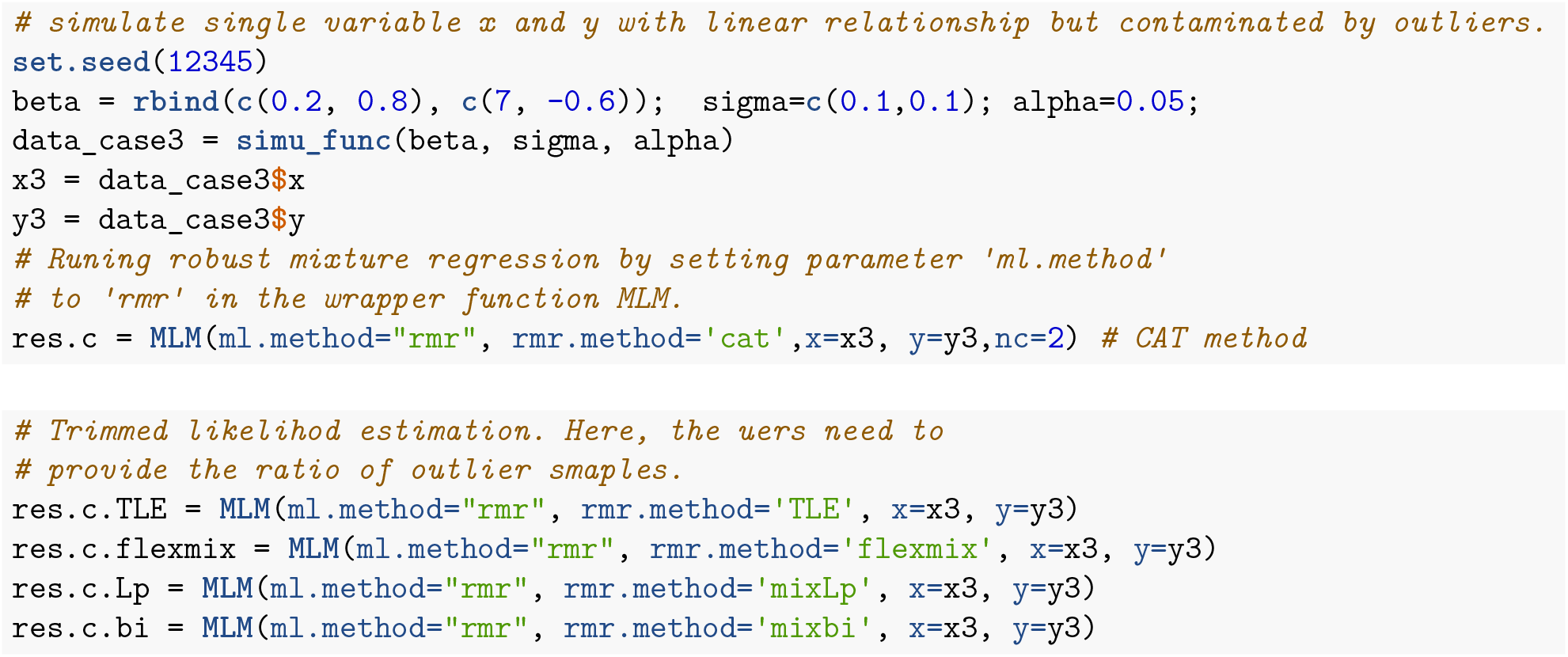

We can extract the outlier samples from the output of MLM.

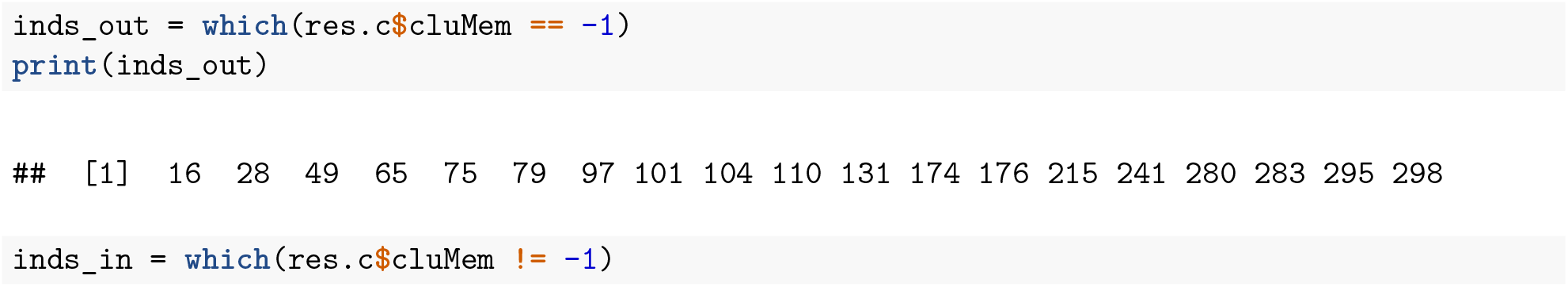

The mixture regression parameters could be extracted by:

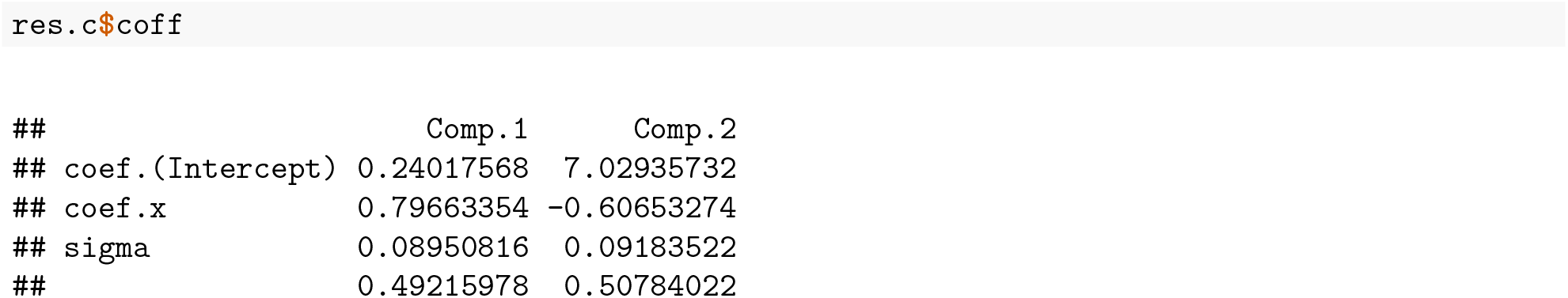

Here, the first two rows are regression coefficients, third row for the standard deviations, and fourth row for the component proportions.

We can visualize the fitted regression and the outliers by ‘compPlot’ function with ‘rlr’ parameter, shows in figure 1.

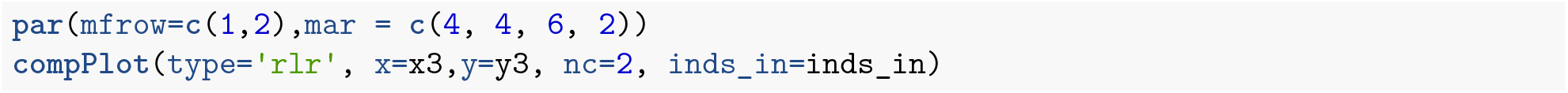

When *K* = 1, this degenerates to ordinary robust linear regression. We call for robust linear regression in the *MLM* wapper function by letting the ‘ml.method’ equal to ‘rlr’. In the core of the method is the ‘ltsReg’ function from the ‘robustbase’ package.

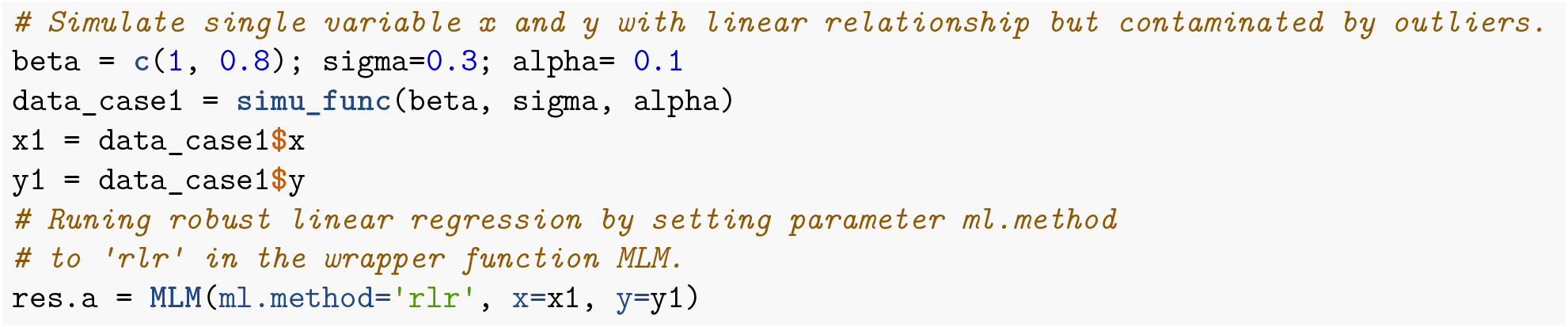

We can extract the outlier samples from the output of MLM.

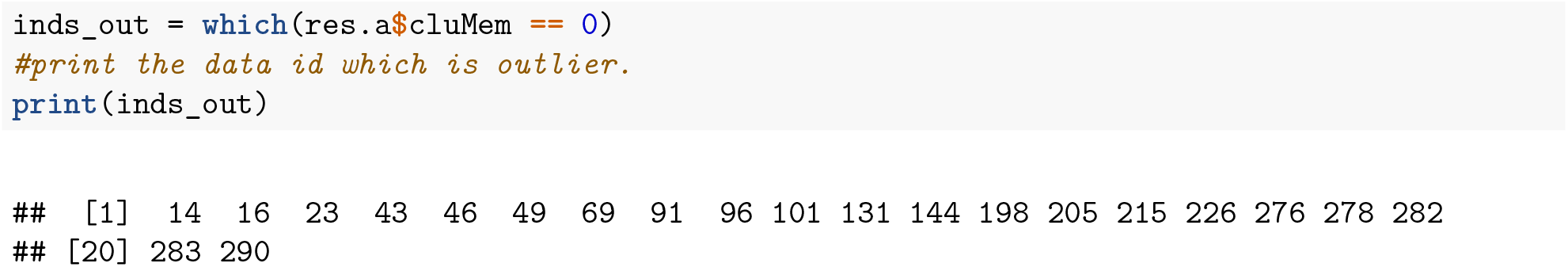

The regression parameters could be extracted by:

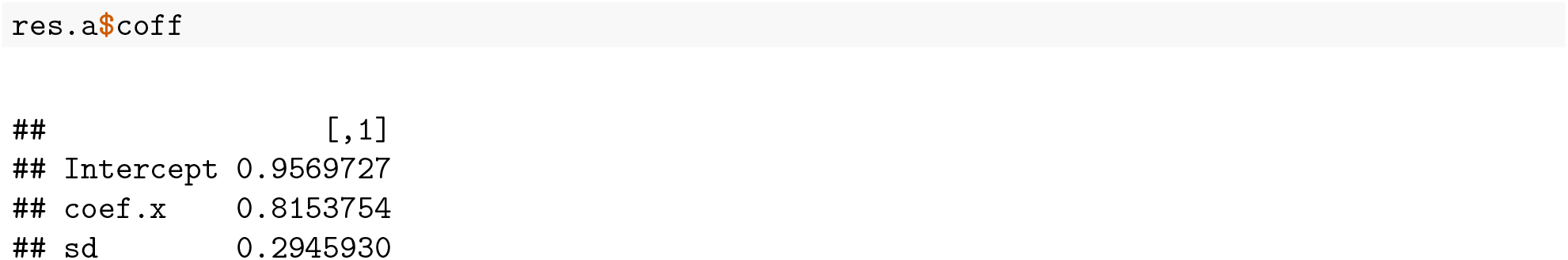

We can visualize the fitted regression and the outliers by *compPlot* function in figure 2.

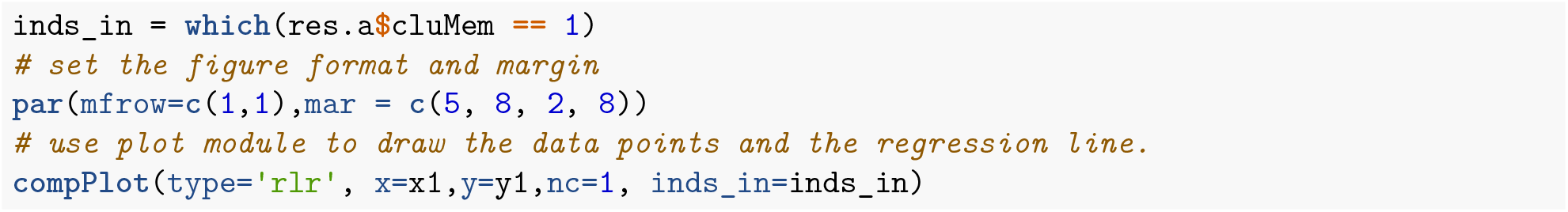

**Figure 2:**
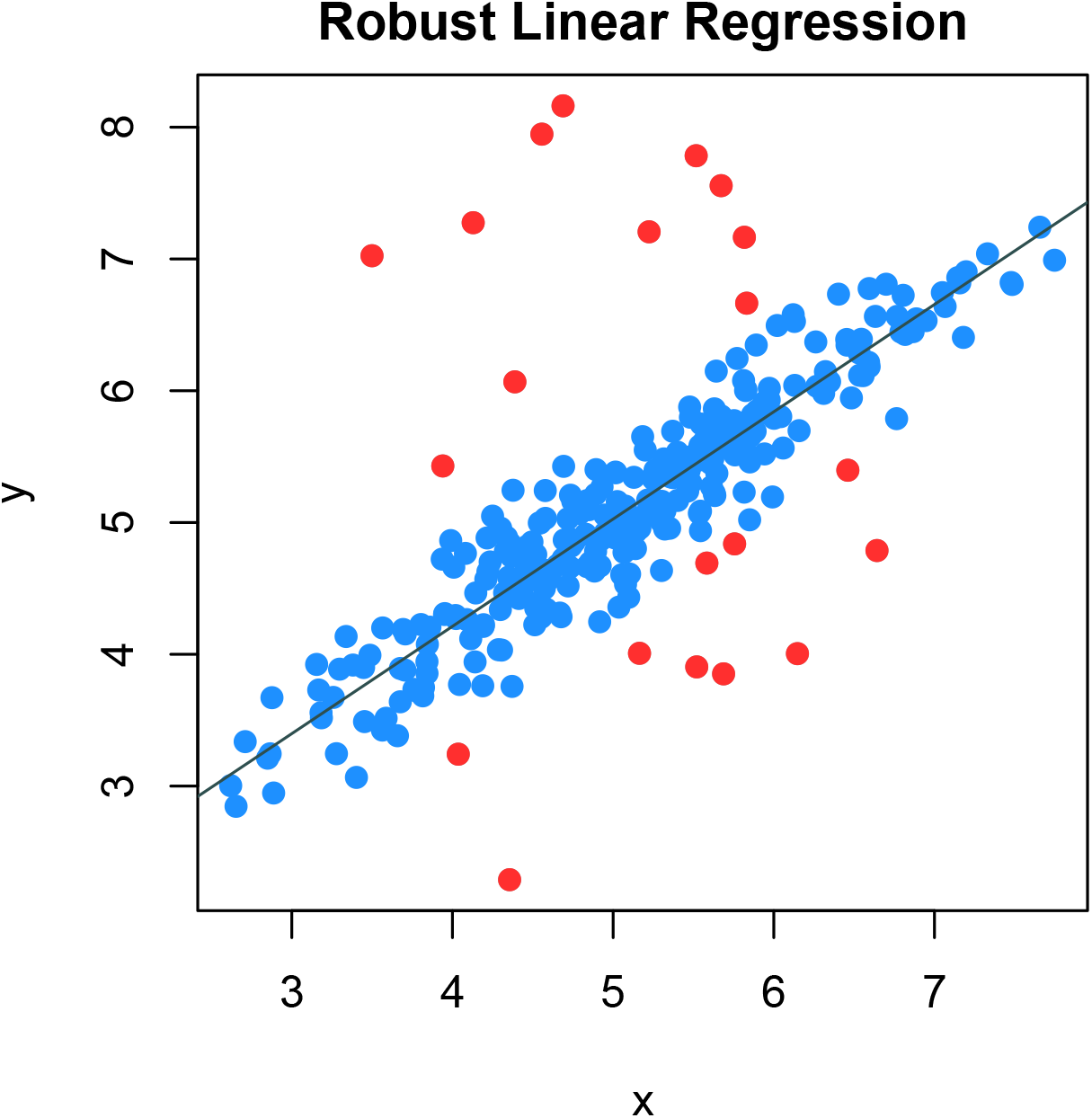
Robust linear regression result

#### S2.2 Mixture regression with flexible modeling

When fitting two or more lines through the data, there could be different regimes with the predictors in different lines. For example, two lines may involve different subsets of the predictors; or a line could include nonlinear transformations of the predictors to relax the “linearity” concerns. Different from the traditional mixture regression, RobMixReg allows for different set of predictors and their transformations as predictors for different component.

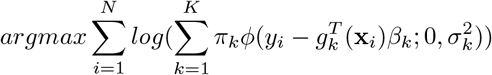

Here, *g*_*k*_(·) is a vectorized function that transforms the original predictors in regression componetn *K*, to combinations of the predictors and their transformations.

RobMixReg can easily achieve flexible modeling by specifying the regression formulas for different components. To enable flexible mixture regression, we call robust mixture regression in the *MLM* wapper function by letting the ‘ml.method’ be ‘fmr’. In the core of the method is the ‘mixtureReg’ function from the ‘mixtureReg’ package.

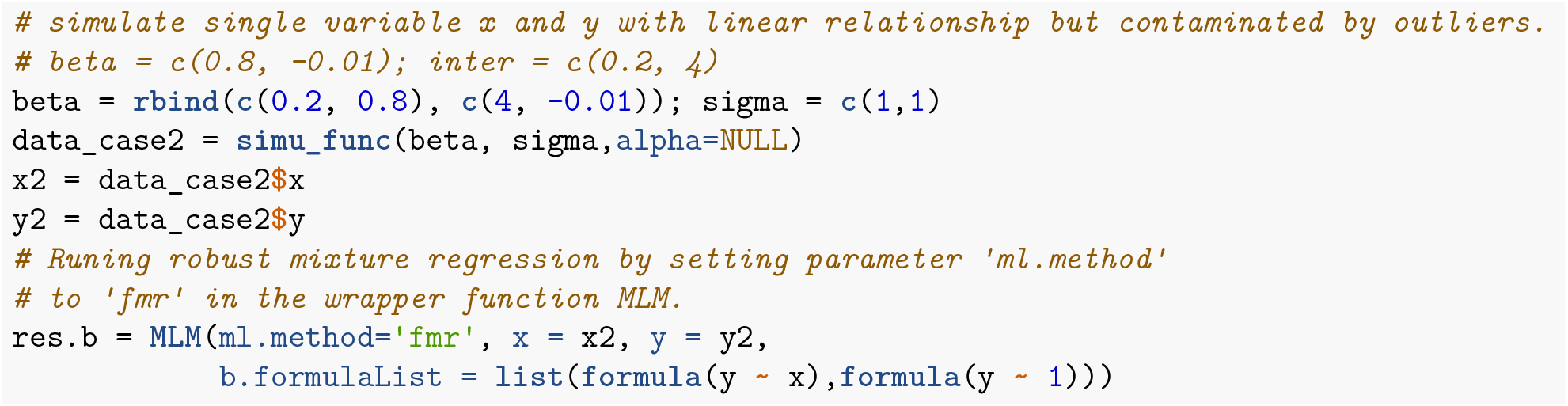

The mixture regression parameters could be extracted by:

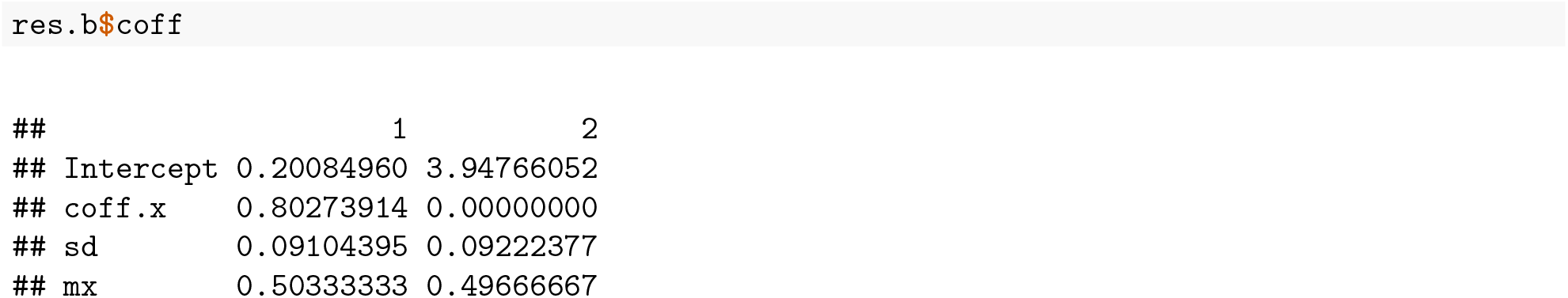

Here, the first two rows are regression coefficients, third row for the standard deviations, and fourth row for the component proportions.

We can visualize the fitted regression lines by *compPlot* function and declare the ‘type’ parameter as ‘mr’ shows in figure 3.

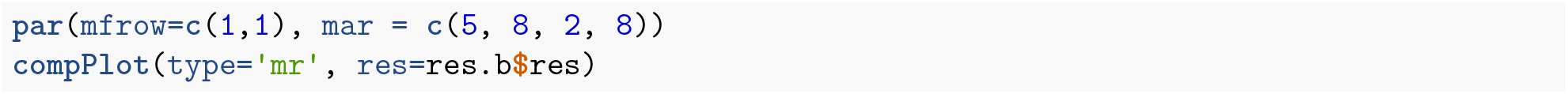

**Figure 3:**
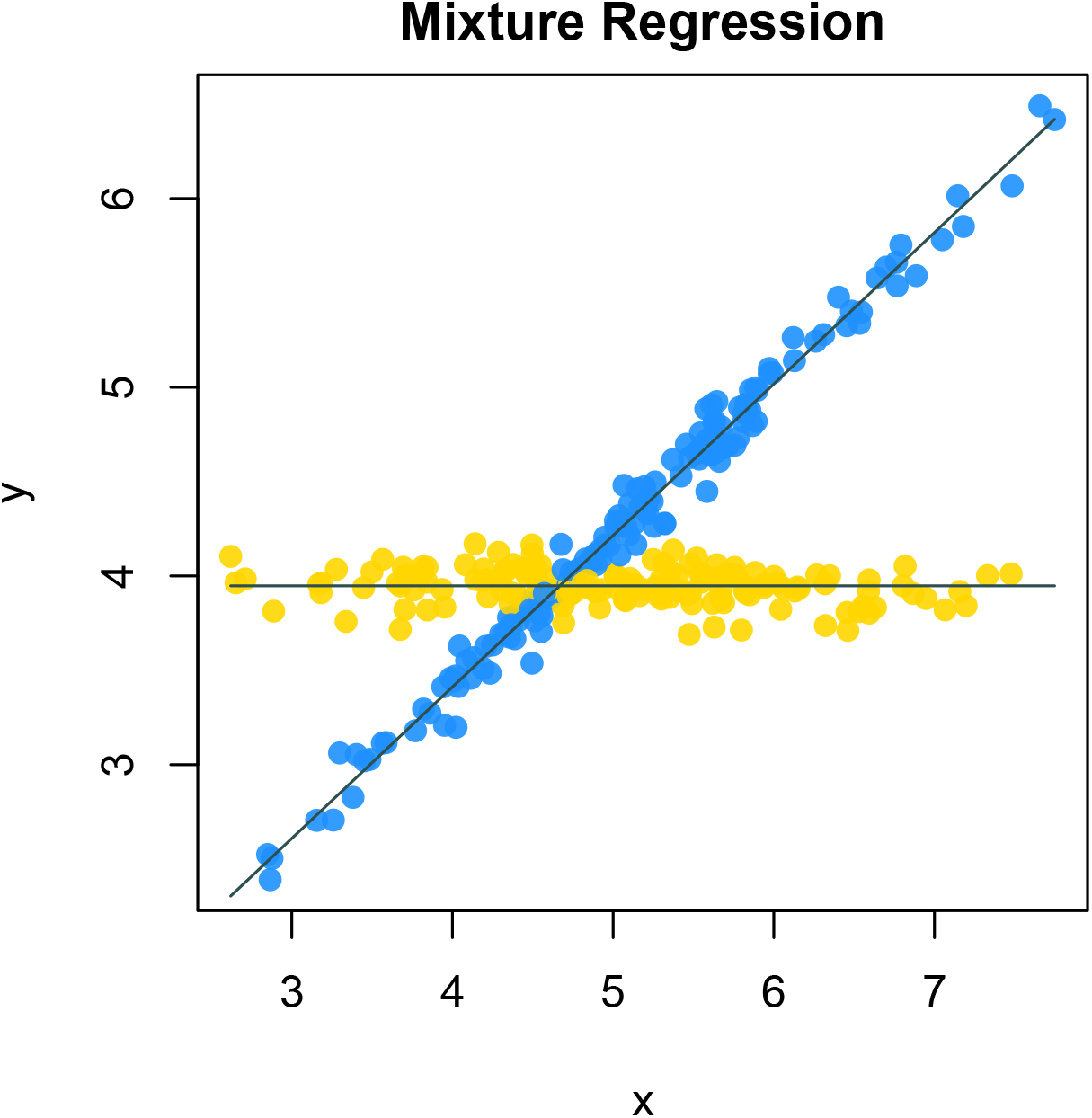
Flexible mixture regression result

#### S2.3 High dimensional mixture regression

Given high dimensional predictors, which is often the case in biological data, we have the total number of parameters to be estimated far more than the total number of observations, making the aofrementioned methods fail. In addition, the dense linear coefficients makes it hard to deduce the subgroup specific features and make meaningful interpreta-tions. We need to add regularization to the regression coefficients to achieve sparse mixture regression.

- Mathmatical Model

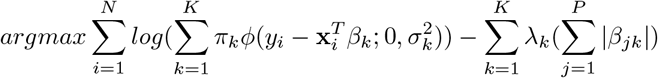

Different from the low dimensional setting, we are imposing constraints on the coefficients through the penalty parameter *λ*_*k*_. The CSMR algorithm is used here, which adaptively selects the penalty parameters *λ*_*k*_, hence there is no need for model selection with regards to *λ*_*k*_.

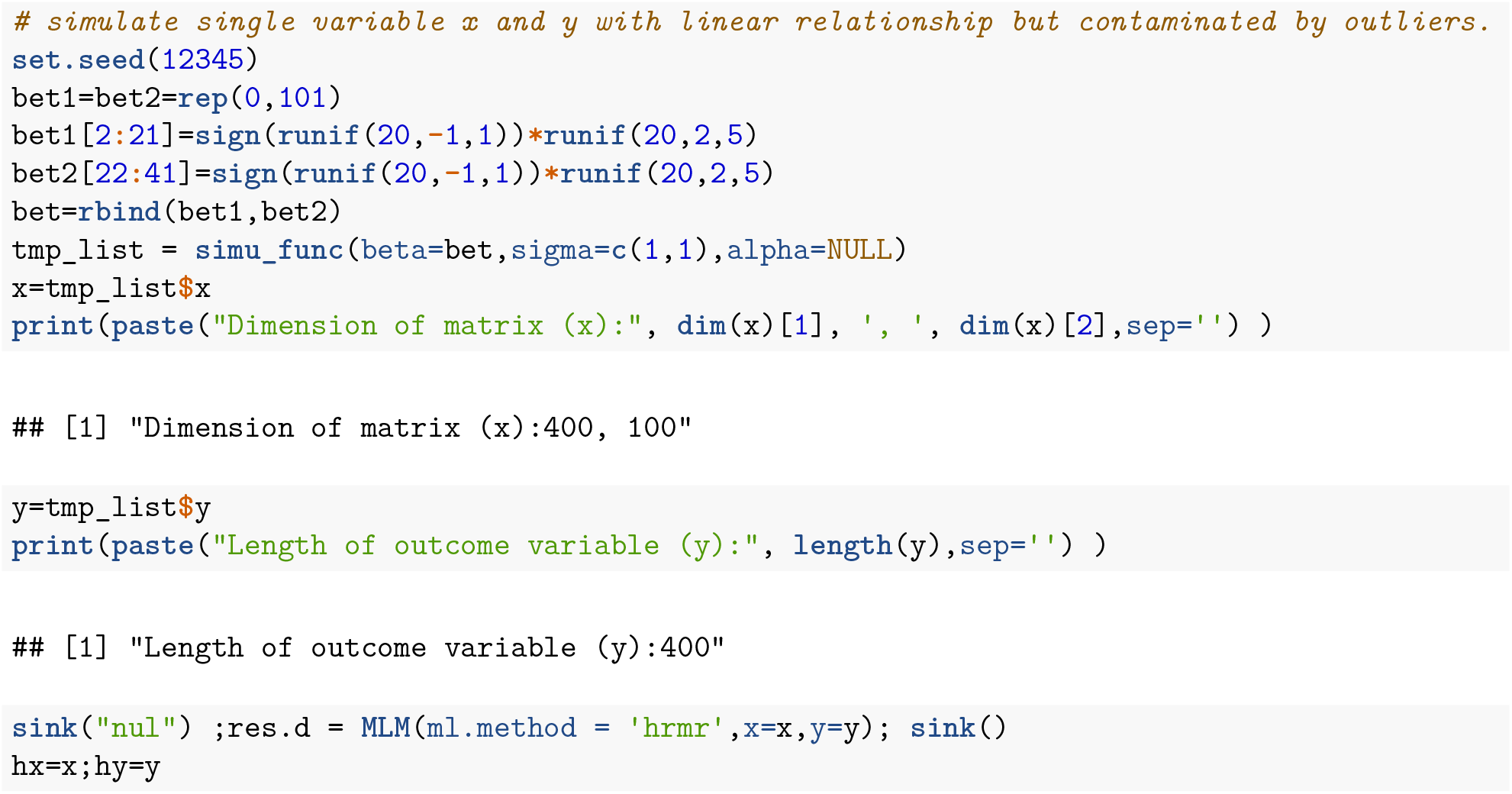

The mixture regression parameters could be extracted by:

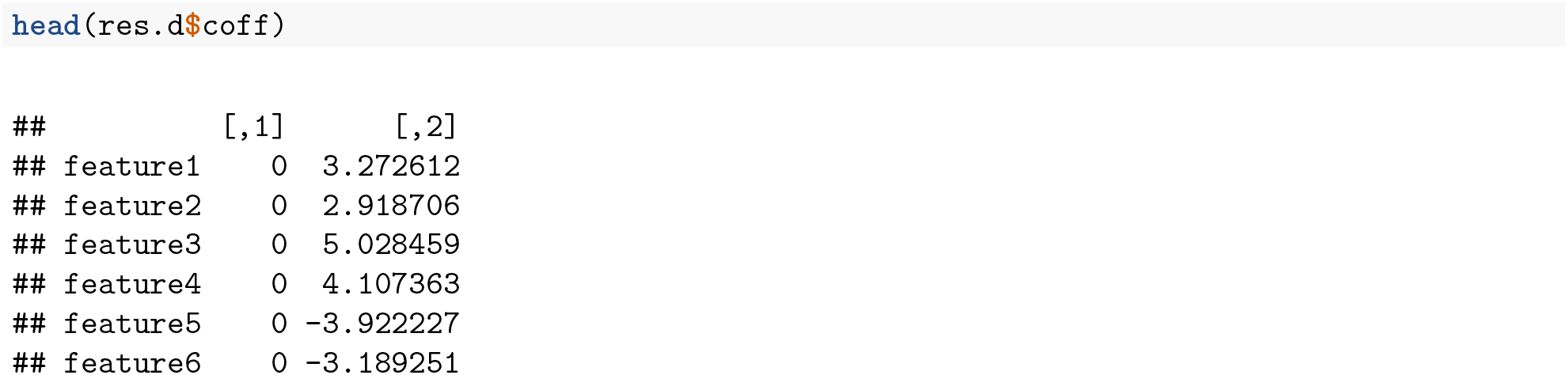

Here, the first (*P* + 1) rows are regression coefficients, (*P* + 2)-th row for the standard deviations, and (*P* + 3)-th row for the component proportions.

We visualize the fitted coefficients using heatmap shows in figure 4. The columns are the clusters and the rows are the features.

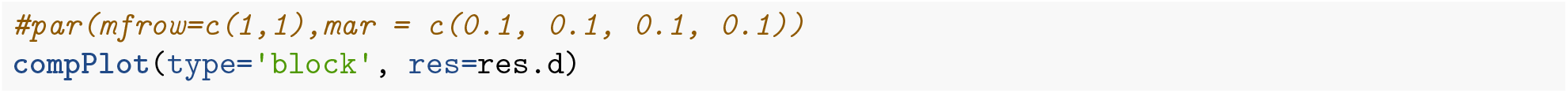

**Figure 4:**
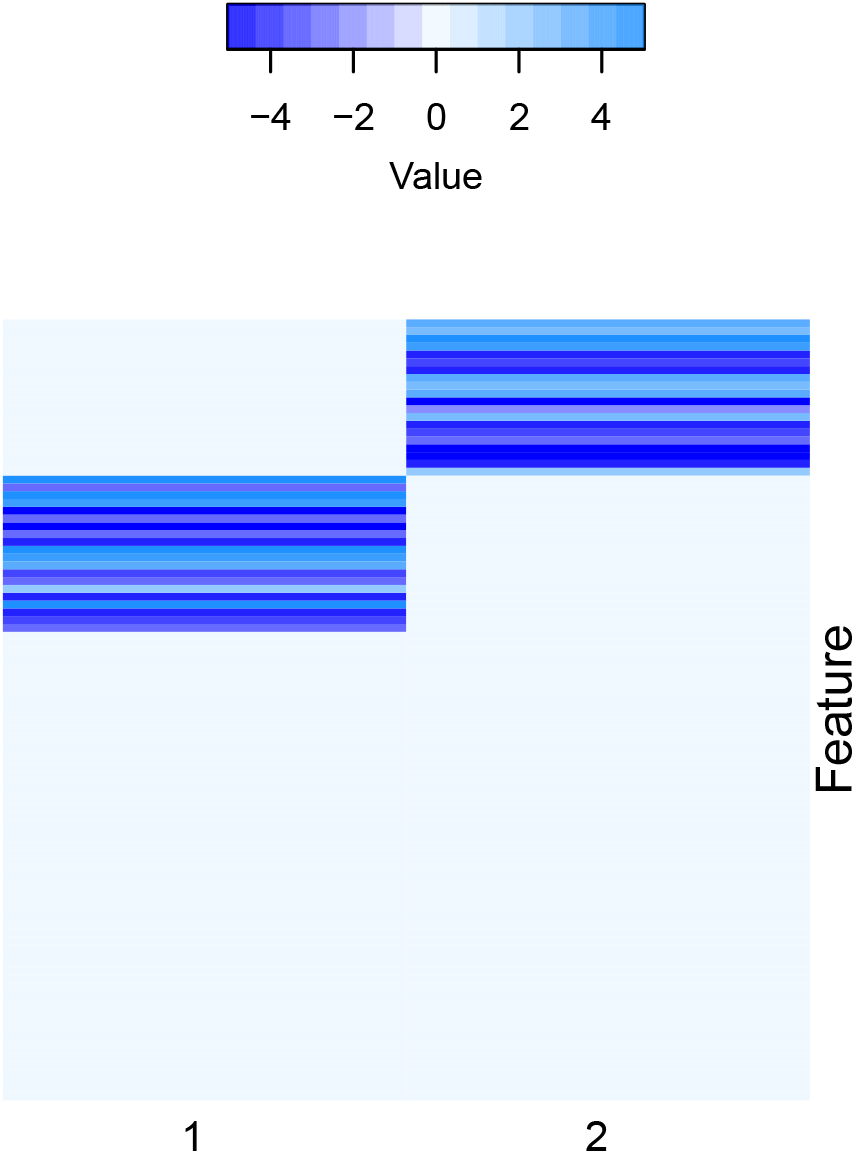
High dimensional feature space robust mixture regression result

#### S2.4 Order Selection

##### BIC based order selection

We provide two methods for selecting the number of components, *K*: Bayesian information criterion (BIC) and cross validation. Meanwhile, in flexible modeling, even with the same *K*, the models could still differ in their complexity, and we also provided BIC metric for model selection. The BIC of each fitted model could be extraced in slot ‘BIC’ by declaring the parameter ‘ml.method’:

Robust mixture regression using BIC

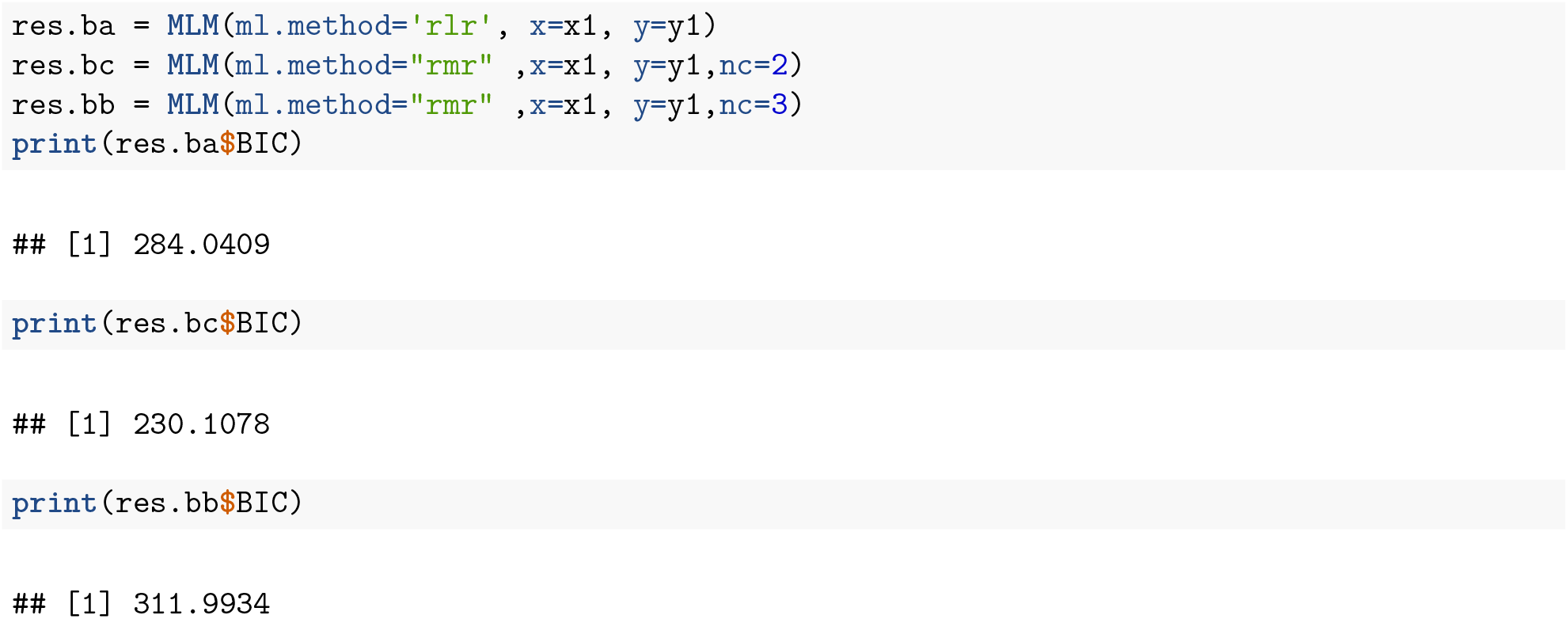

Flexbile mixture regression using BIC

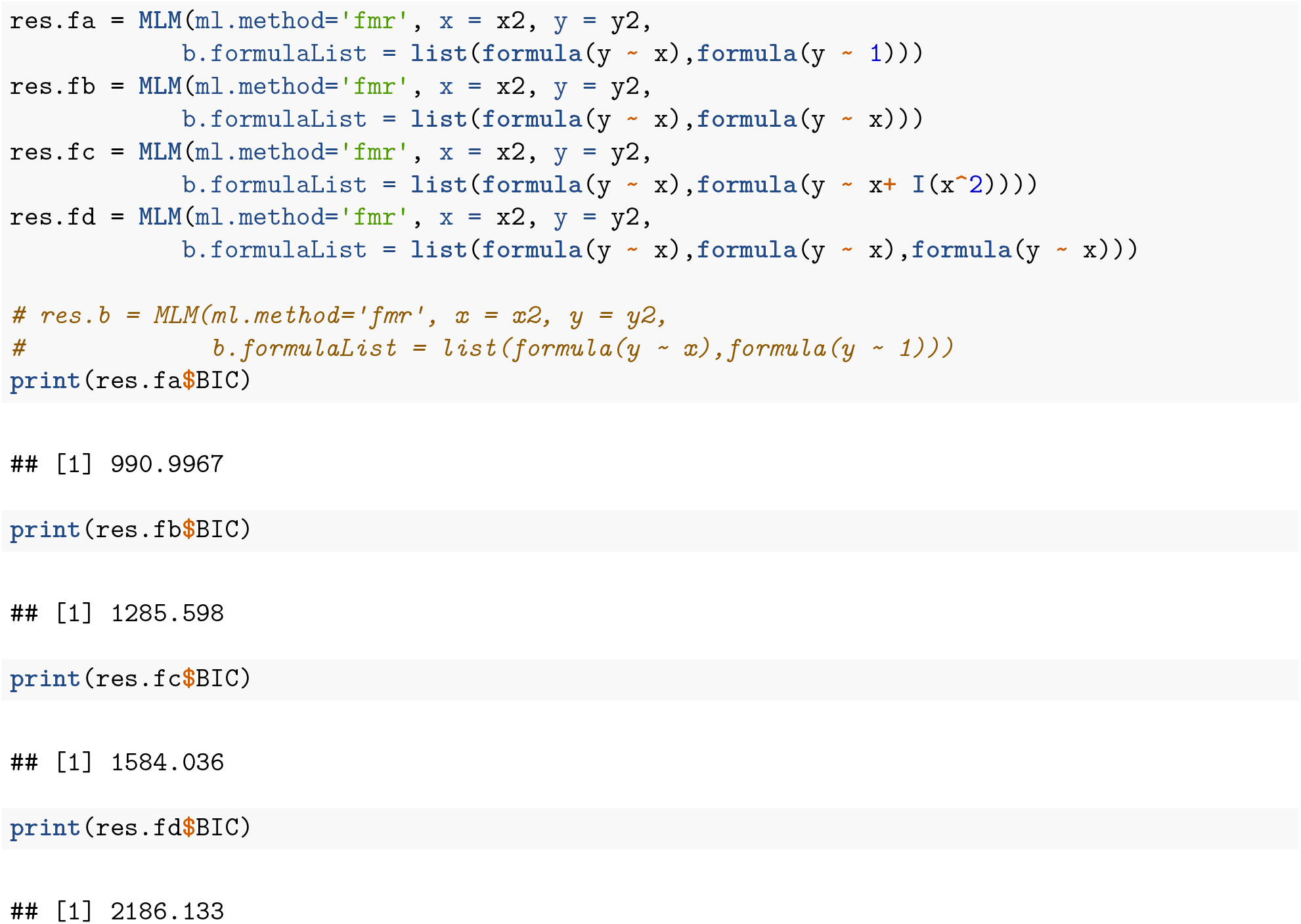

High dimensional mixture regression using BIC

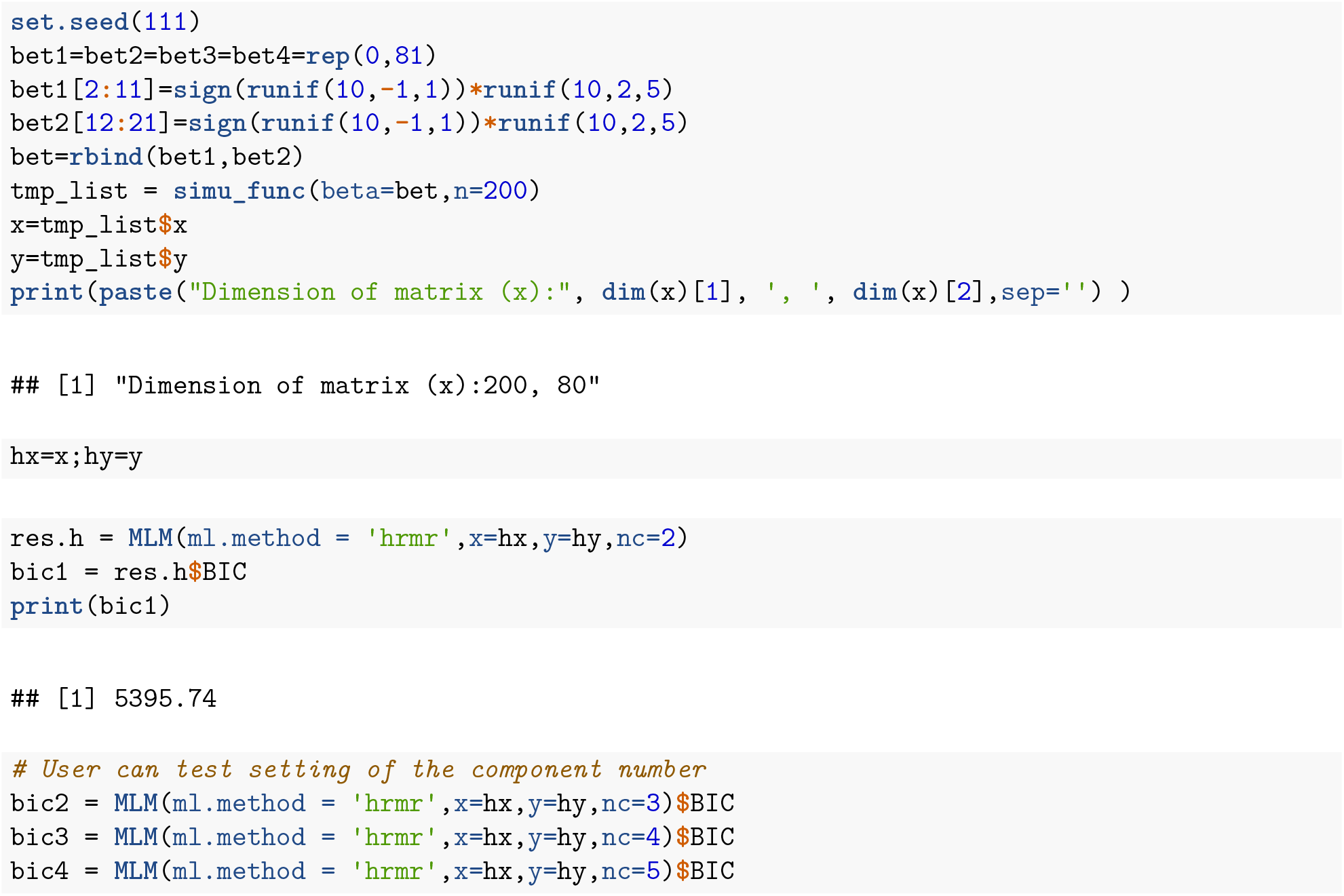

##### Cross validation based order selection

For high dimensional predictors, we also provide cross validation for selection of *K*.A large *K* will tend to overfit the data with more complex model of higher variance, while smaller *K* might select a simpler model with larger bias. Take a 5-fold cross validation as an example. For given *K*, at each repetition, 80% samples are used for training to obtain the mixture regression parameters. Then, for any sample from the rest of the testing data, it is first assigned to a cluster by maximum posterior probability; then a prediction of 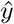 could be made based on the model parameters given *x*. Using the testing data, we could decide how to balance the trade-off between bias and variance. To evaluate how the estimated model under *K* explains the testing data, we could calculate the root-meansquare-error between the observed *y* and the predicted 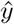, or Pearson correlation between the two. By repeating this procedure for multiple times, a more robust and stable evaluation of the choice of *K* should be derived based on the summarized RMSE or Pearson correlations.

The output of cross validation function, *MLM_cv*, is the average correlation or RMSE on the testing data. Higher correlation and lower RMSE indicate a better model with good bias variance tradeoff.

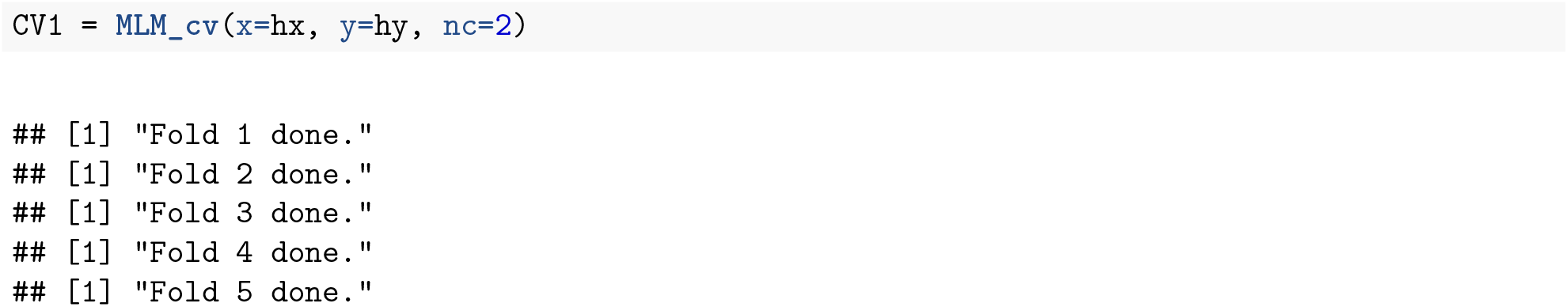

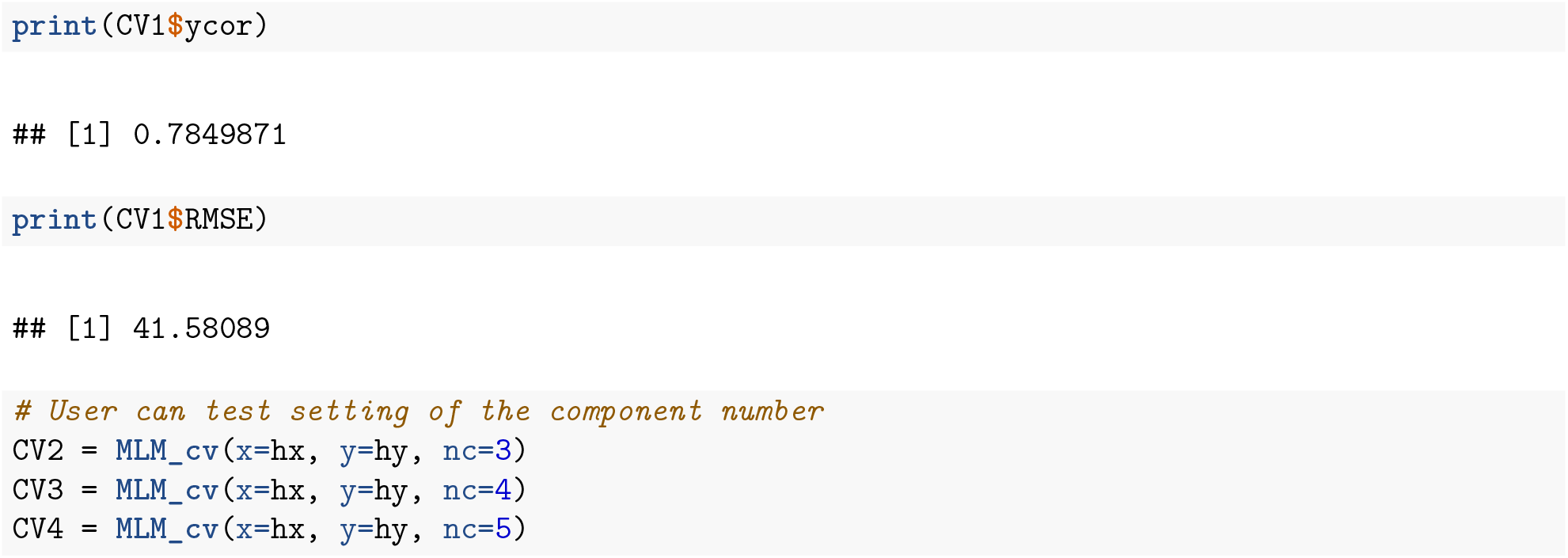

### S3. Real Data Application

#### S3.1 Colon adenocarcinoma disease application

Colon adenocarcinoma is known as a heterogeneous disease with different molecular subtypes. We demonstrate the usage of o*RobMixReg*on collected expression of a few genes and the methylation profiles for some of their CpG sites from the Cancer Genome Atlas (TCGA) cohort.

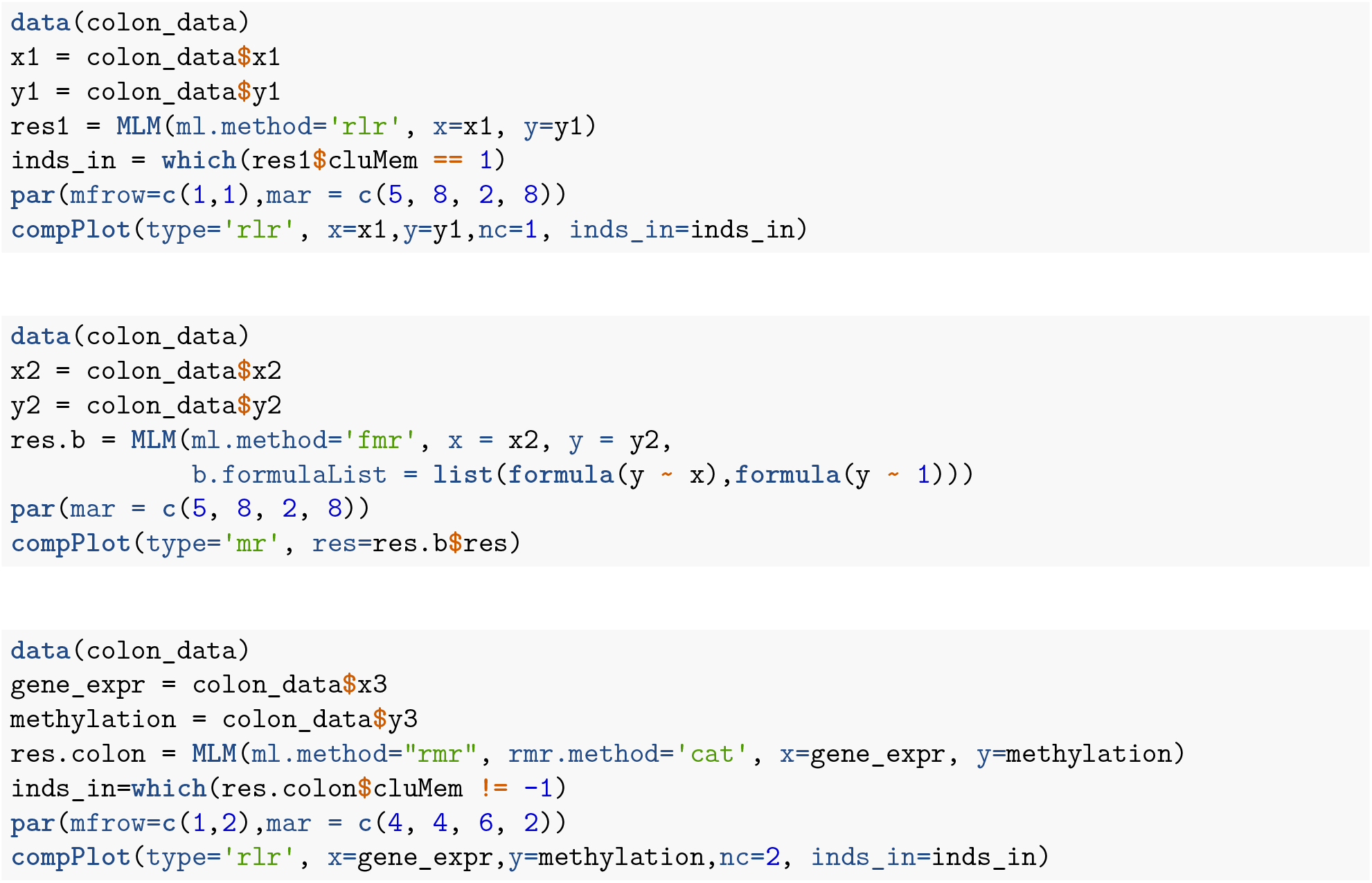

We fitted the data using CAT, the two colored line represented the two subgroups of patients.

**Figure 5:**
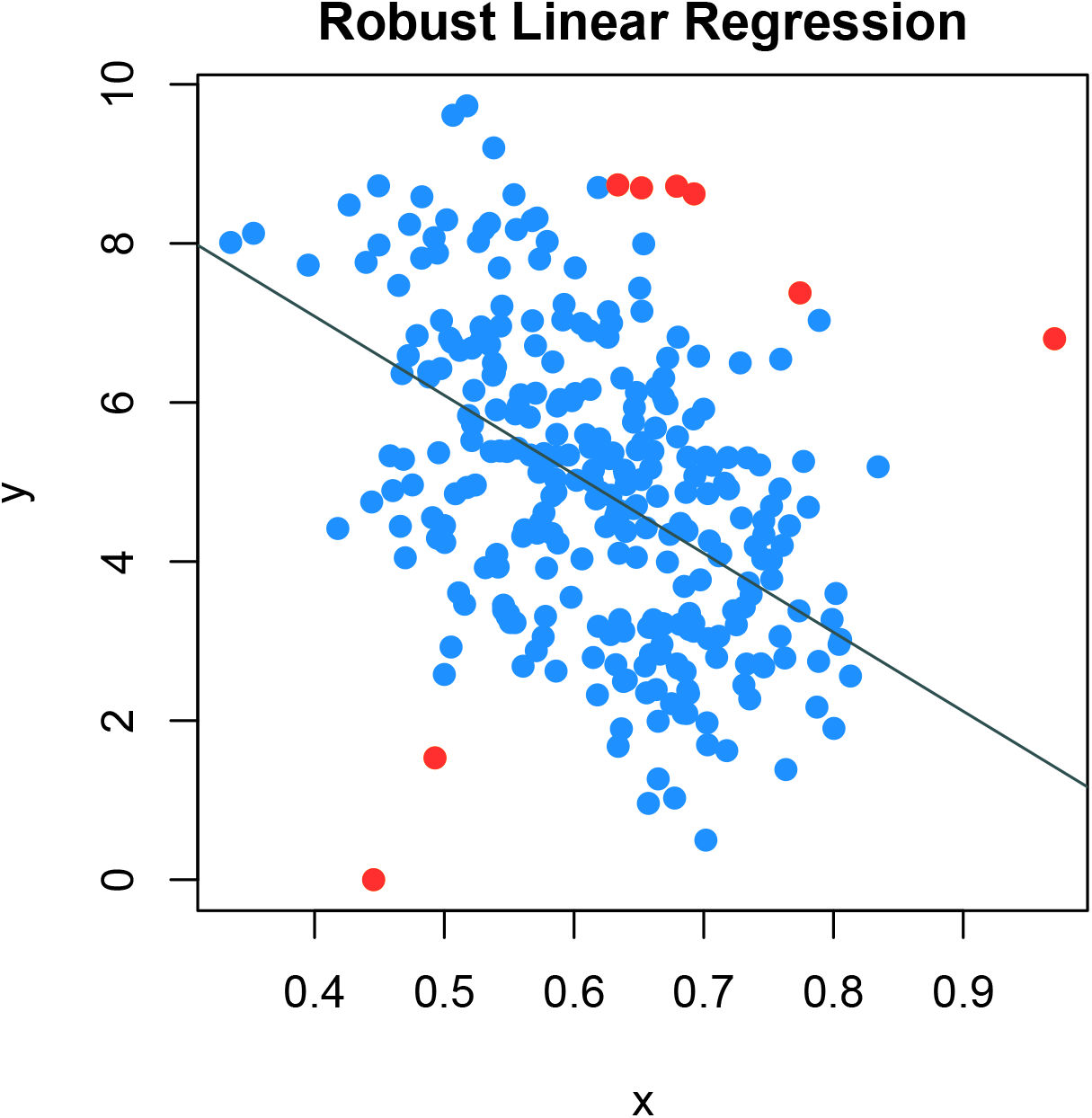
Colon data robust linear regression

**Figure 6:**
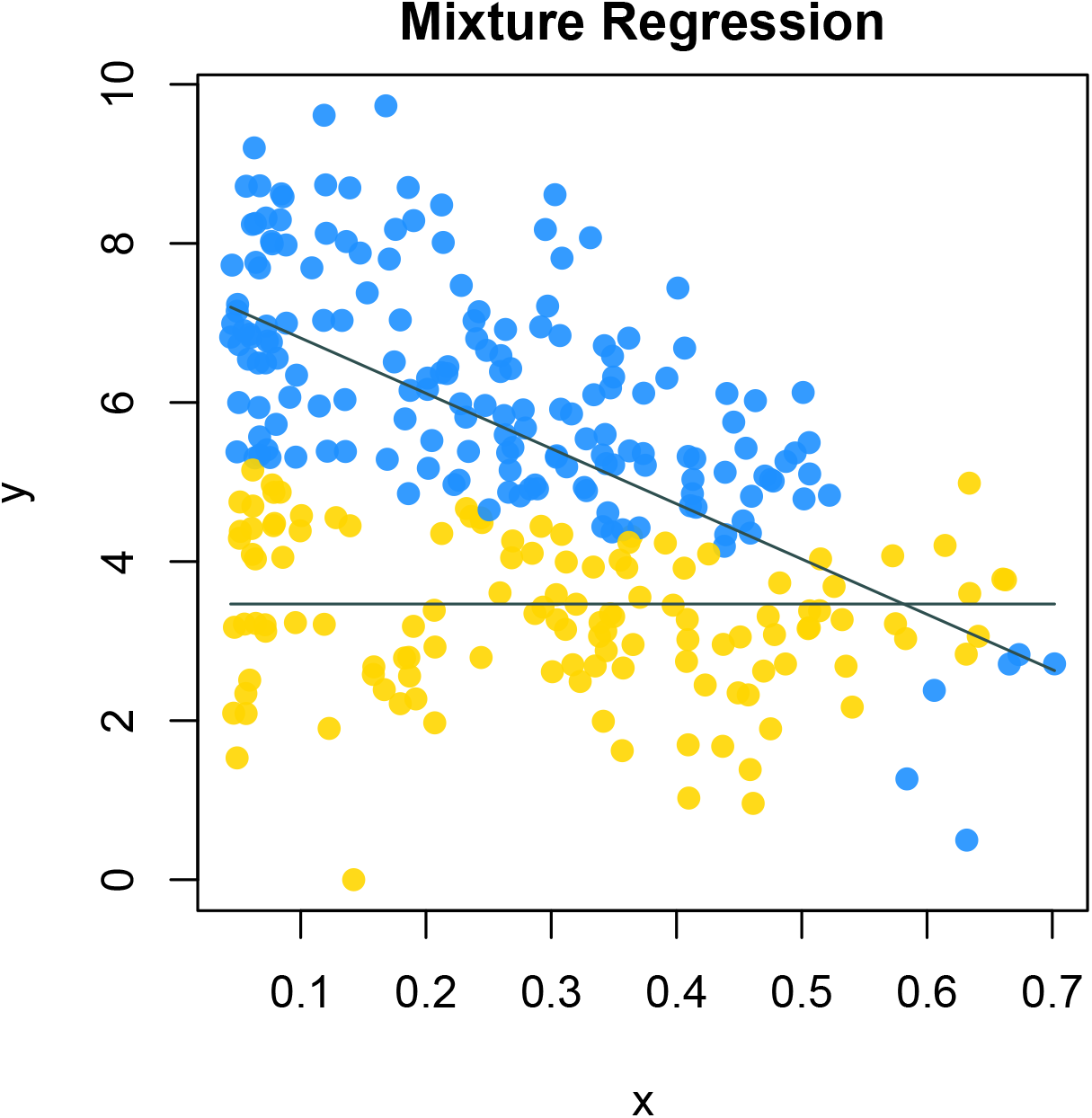
Colon cancer data flexible mixture regression

**Figure 7:**
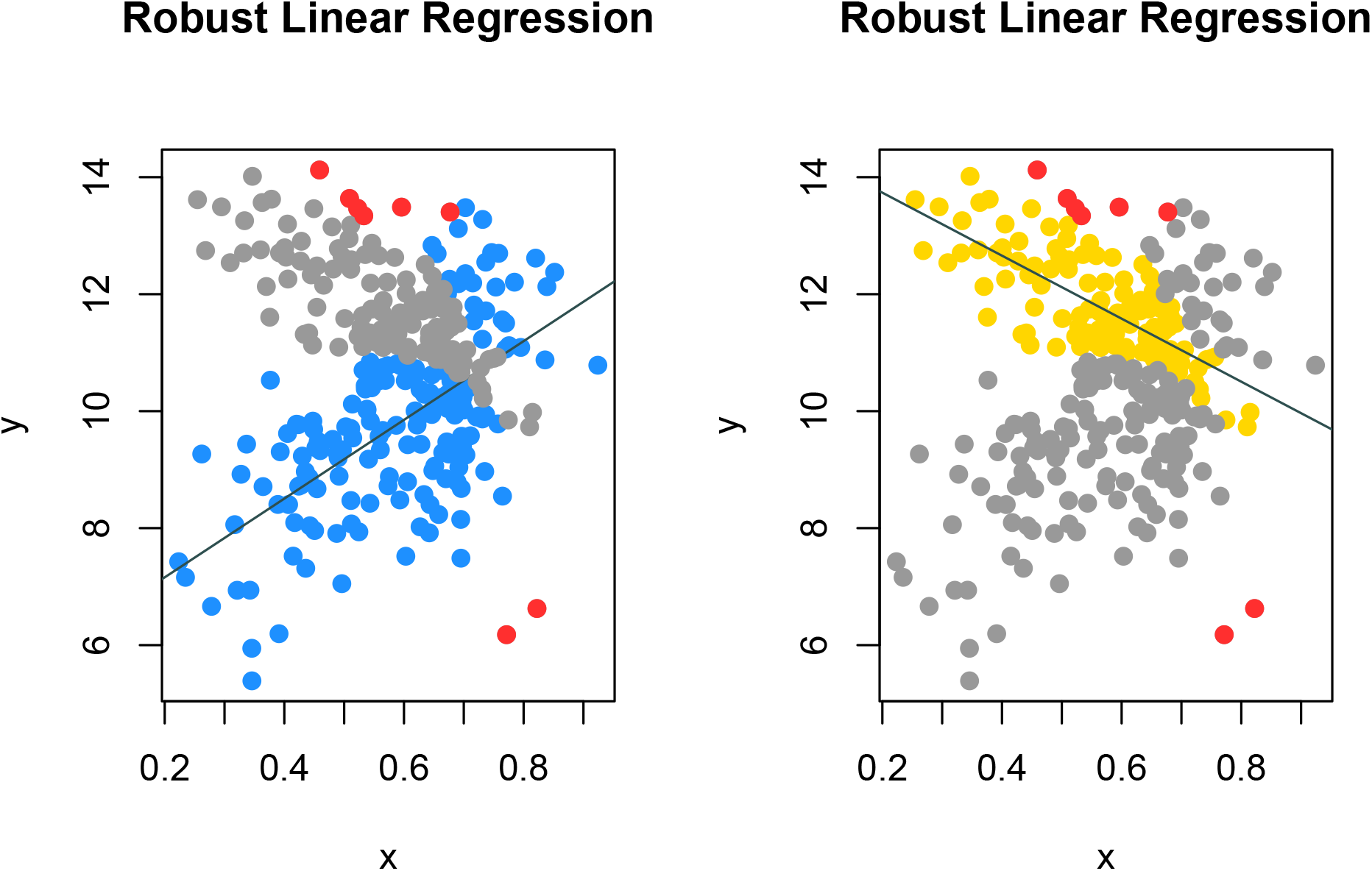
Colon cancer data robust mixture regression

#### S3.2 Subspace clustering on high dimension dataset - CCLE application

We collected gene expression data of 470 cell lines as well as the cell lines’ sensitivity score (AUCC score) for ‘AEW541’ drug from the Cancer Cell Line Encyclopedia (CCLE) dataset. We demonstrate the usage of our package on this high dimensioanl dataset.

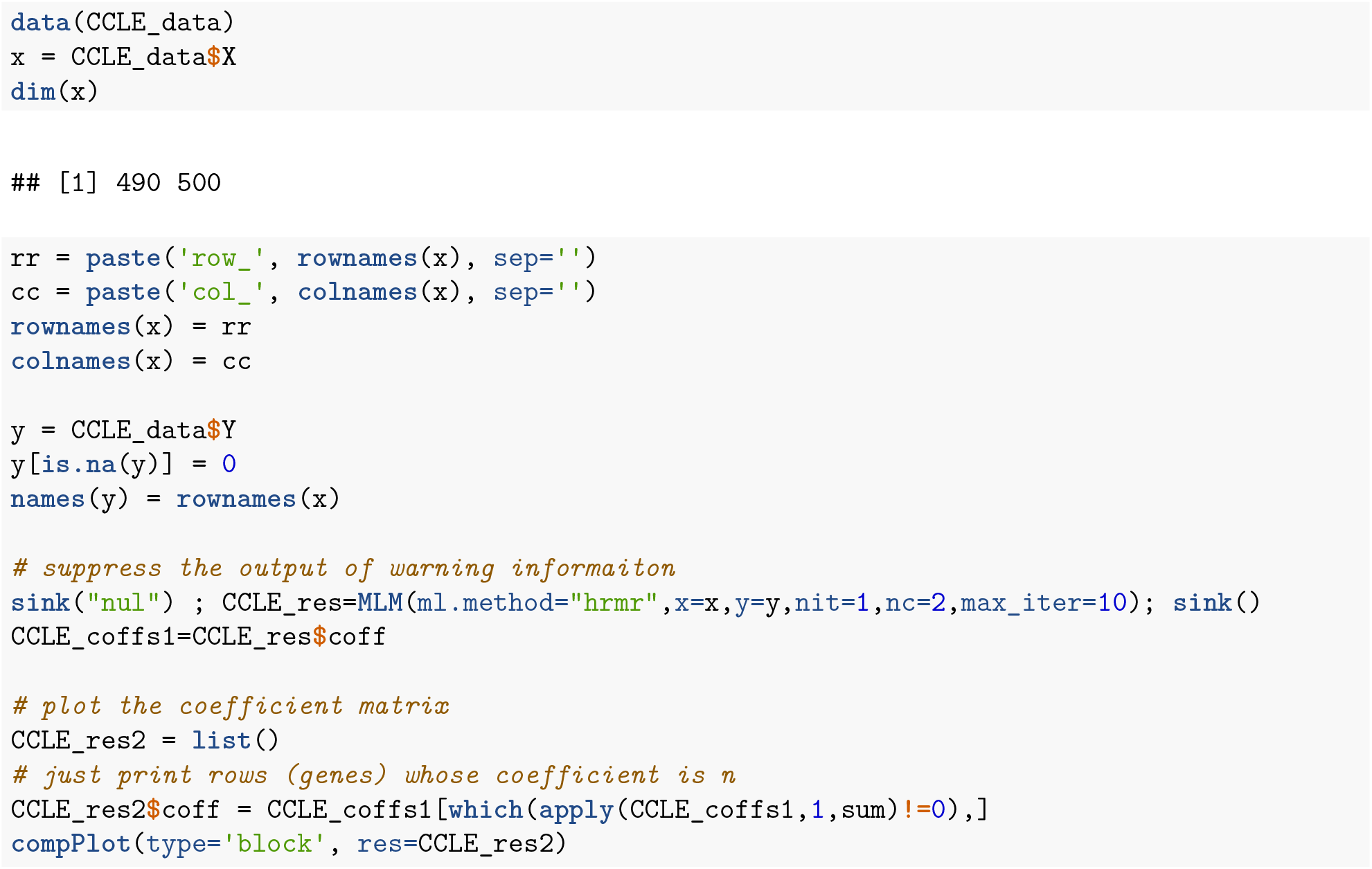

**Figure 8:**
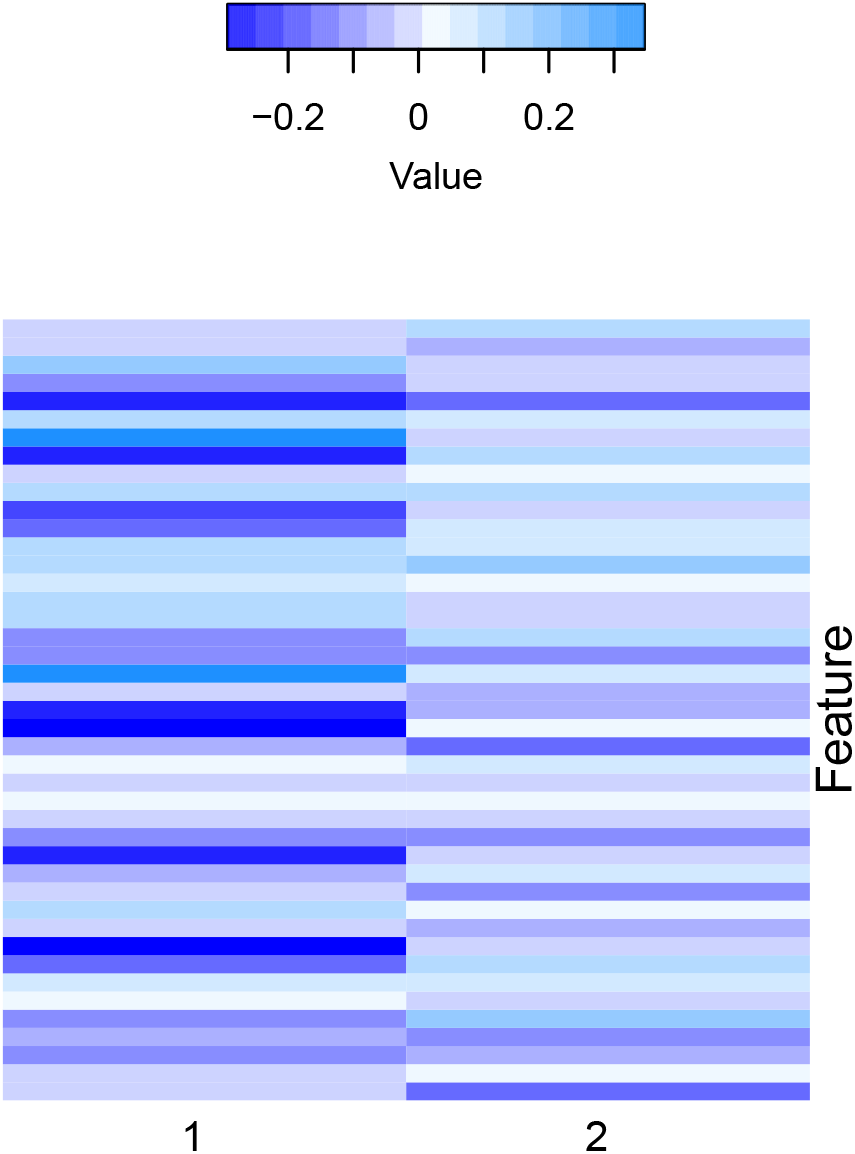
High dimension space feature robust mixture regression result

